# Effective targeting of CD19 positive primary B-ALL cells using CAR-NK cells generated with mRNA-LNPs

**DOI:** 10.1101/2024.11.21.624707

**Authors:** Hossein Salehi-Shadkami, Mosslim Sedghi, Shima Tavoosi, Masoumeh Alimohammadi, Reza Alimohammadi, Maryam Barkhordar, Ahmadreza Mofayezi, Mohammad Sadra Modaresi, Mohammad Vaezi, Somaye Dehghanizadeh, Mohammad Ahmadvand, Vahid Khoddami

## Abstract

Chimeric antigen receptor natural killer (CAR-NK) cell therapy is recognized as a promising modality for the treatment of hematologic malignancies, particularly B-cell malignancies. In this study, we developed “off-the-shelf” anti-CD19 CAR-NK cells using anti-CD19 CAR mRNAs formulated in proprietary ionizable lipid nanoparticles (LNPs). The efficiency of mRNA-LNP delivery into umbilical cord blood (UCB)-derived NK cells and primary T cells was evaluated in an in-vitro setting, demonstrating superior delivery efficiency in NK cells. Further investigation showed a probable role for an endocytic mechanism, macropinocytosis, in efficient transfection of NK cells with LNPs. Nevertheless, CAR-NK cells generated through this mRNA-LNP platform exhibited significantly enhanced cytotoxicity against CD19^+^ target cells, such as EGFP^+^Raji stable cell line and primary malignant B cells derived from refractory/relapsed B-cell acute lymphoblastic leukemia (B-ALL) patients. These findings highlight the promise of the mRNA-LNP platform in advancing CAR-NK therapies against B-cell malignancies.

## 2. Introduction

B-cell malignancies, including B-cell leukemias (chronic lymphocytic leukemia [CLL] and B-ALL), B-cell lymphomas (the most common type of non-Hodgkin lymphoma [NHL]), and plasma cell dyscrasia, constitute a significant portion of hematological cancers with distinct epidemiological patterns. While NHL and CLL predominantly affect older individuals, B-ALL demonstrates a bimodal age distribution peaking in early childhood and young adulthood. Despite recent advancements in treatments, particularly for pediatric B-ALL, B-cell malignancies remain a significant cause of morbidity and mortality, underscoring the need for continued research and comprehensive management strategies to address the burden of morbidity and mortality across all age groups [1–4].

In recent years, CAR-armored cell therapy has gained increasing attention, particularly in the treatment of hematological malignancies [5, 6] with a special focus on T cells [7]. CAR-T cells have shown promise as an effective therapeutic approach for combating various tumor types [8]. To date, seven CAR-T cell-based drugs have received approval from the FDA for the treatment of hematological malignancies [9, 10], and promising clinical results have been reported for children and young adults with relapsed/refractory B-ALL in CAR-T cell therapy clinical trials [11–17]. While effective CAR-T cell therapies hold significant promise, they face several challenges in the clinical setting. These include high treatment costs per patient and a considerable incidence of toxicities, such as cytokine release syndrome (CRS), neurotoxicity, and graft-versus-host disease (GvHD) [18–21]. Current research is focused on developing strategies to address challenges associated with CAR-T cell therapies in clinical practice [22, 23].

Utilizing the power of CAR-armored cell therapy, ongoing clinical trials are investigating various immune cell subsets, such as CAR-NK, CAR-Monocyte, CAR-Treg, and CAR-NKT cell therapies [24–31]. Among these, CAR-NK cell therapy, with its ‘‘off-the-shelf nature’’, allows for the administration of allogeneic cells with reduced risks of CRS and GvHD [32–36]. In fact, CAR-NK cell therapy has shown great promise in treating not only hematological malignancies such as B-ALL, diffuse large B-cell lymphoma (DLBCL), NHL, acute myeloid leukemia (AML), and multiple myeloma; but also solid tumors, including breast cancer, lung cancer, glioblastoma, renal cancer, hepatocellular carcinoma, pancreatic cancer, and gastric cancer [36, 37].

The sources of NK cells for CAR-NK cell therapies include NK cell lines such as NK-92, NK-92MI, and KHYG-1, as well as peripheral blood NK cells, placental NK cells, UCB-derived NK cells, and stem cell-derived NK cells. Each source presents its own advantages and limitations [38, 39]. Despite concerns regarding UCB-NK cell use, such as immature phenotype and relatively lower cytotoxicity compared with other NK cell sources, they offer several advantages, including ease of accessibility in UCB bank, large expansion base, and better proliferation capacity.

In a recent Phase I/II trial, 37 patients with refractory/relapsed B-cell malignancies were treated with CD19-targeted, UCB-derived CAR-NK cells, generated with retrovirus vector. Results showed an overall response rate of 48.6% at 100 days post-treatment, with overall survival rates and 1-year progression-free of 68% and 32%, respectively. Notably, the trial observed a 100% response rate in low-grade non-Hodgkin lymphoma at 30 days and significant responses in chronic lymphocytic leukemia and diffuse large B-cell lymphoma. Key factors for optimal outcomes included using UCB units cryopreserved within 24-hours and using best-performing units with a low nucleated red blood cell content. The treatment demonstrated a well-tolerated safety profile, with no severe adverse events reported [40, 41].

Viral vectors, including retroviruses and lentiviruses, are commonly employed for the ex vivo engineering of NK cells in clinical trials [33, 42–44]. mRNA-LNP technology, on the other hand, offers a safe and efficient alternative for CAR expression inside NK cells [45–49]. This technology not only provides the feasibility of transfection with higher expression rates in NK cells but also mitigates concerns associated with viral vectors such as oncogenicity and immunogenicity [47, 48, 50, 51].

For CAR mRNA transfer into T or NK cells both electroporation and LNP carriers can be used. While electroporation may initially achieve higher CAR expression levels, it is quite often associated with global gene expression perturbation, faster mRNA degradation, and increased cell exhaustion. In contrast, LNPs facilitate longer-lasting CAR expression due to the enhanced persistence of CAR mRNA in transfected cells, resulting in sustained functionality and less cellular damage compared with electroporation. This makes mRNA-LNPs a promising modality for CAR-armored cell therapies[52].

We have previously established our mRNA-LNP platform by developing Covid-19 mRNA vaccine which was successfully evaluated in both preclinical and clinical settings [53, 54]. Here in this study, we extended the applications of our platform, with some modifications, for the generation of CAR-NK and CAR-T cells. We investigated the transfection efficiency of our proprietary LNPs for mRNA delivery in UCB NK cells and T cells derived from PBMCs of healthy donors. We also studied the possible role of macropinocytosis as a key step in NK cell LNP transfection. Based on the outcomes, we selected NK cells for CAR-NK cell generation as the effector cells for subsequent functional assays. We then evaluated CAR-NK cytotoxicity against the EGFP^+^Raji cell line, representing NHL and CD19^+^ B-cell malignancies, and primary B-ALL cells derived from refractory/relapsed B-ALL patients.

## 3. Results

### 3.1. Transfection efficiency of mRNA-LNPs in UCB NK and T cells

Encapsulation of mRNA encoding Enhanced Green Fluorescent Protein (EGFP) and anti-CD19 CAR was achieved using LNPs via a microfluidic technology protocol. Characterization of the final product of LNPs revealed an average size of 109.2 nm with a polydispersity index (PDI) of 0.130 for EGFP mRNA-LNPs, and an average size of 117.1 nm with a PDI of 0.188 for anti-CD19 CAR mRNA-LNPs (Fig.1).

**Fig. 1:**
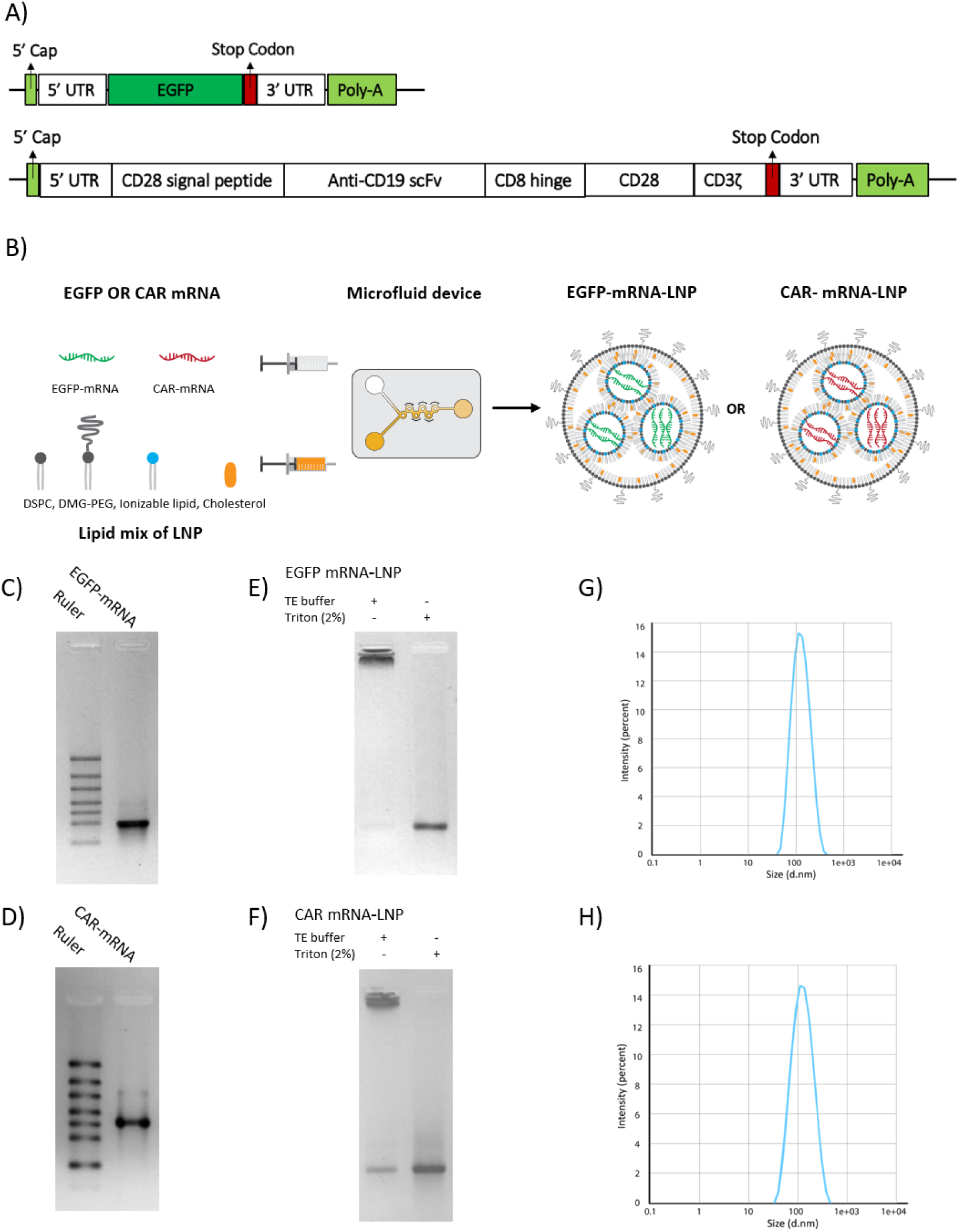
mRNA design, production, and encapsulation in LNPs. A) EGFP and Anti-CD19 mRNA Structures B) schematic illustration of EGFP and CAR mRNA-LNP production. Evaluation of the integrity and purity of the produced EGFP mRNA (C) and anti-CD19 CAR mRNA (D) with gel electrophoresis. Evaluation of EGFP mRNA (E) and CAR mRNA (F) encapsulation efficiency in LNP by Gel Retardation Assay. Evaluation of the size distribution of EGFP mRNA-LNP (G) and anti-CD19 CAR mRNA-LNPs (H) with Dynamic Light Scattering (DLS).

We sought to evaluate our proprietary LNP transfection efficiency in NK and T cells (Fig.2 A). NK cells were isolated from UCB and expanded from day 0 to day 7. To evaluate the transfection efficiency of LNPs in UCB NK cells, 1-2 μg EGFP mRNA-LNPs per 10^6^ NK cell on day 7 were administered. Flow cytometry analysis was conducted to quantify the percentage of EGFP-positive NK cells, revealing transfection efficiencies of 94.7%, 24-hour post-transfection (Fig.2 B).

**Fig. 2:**
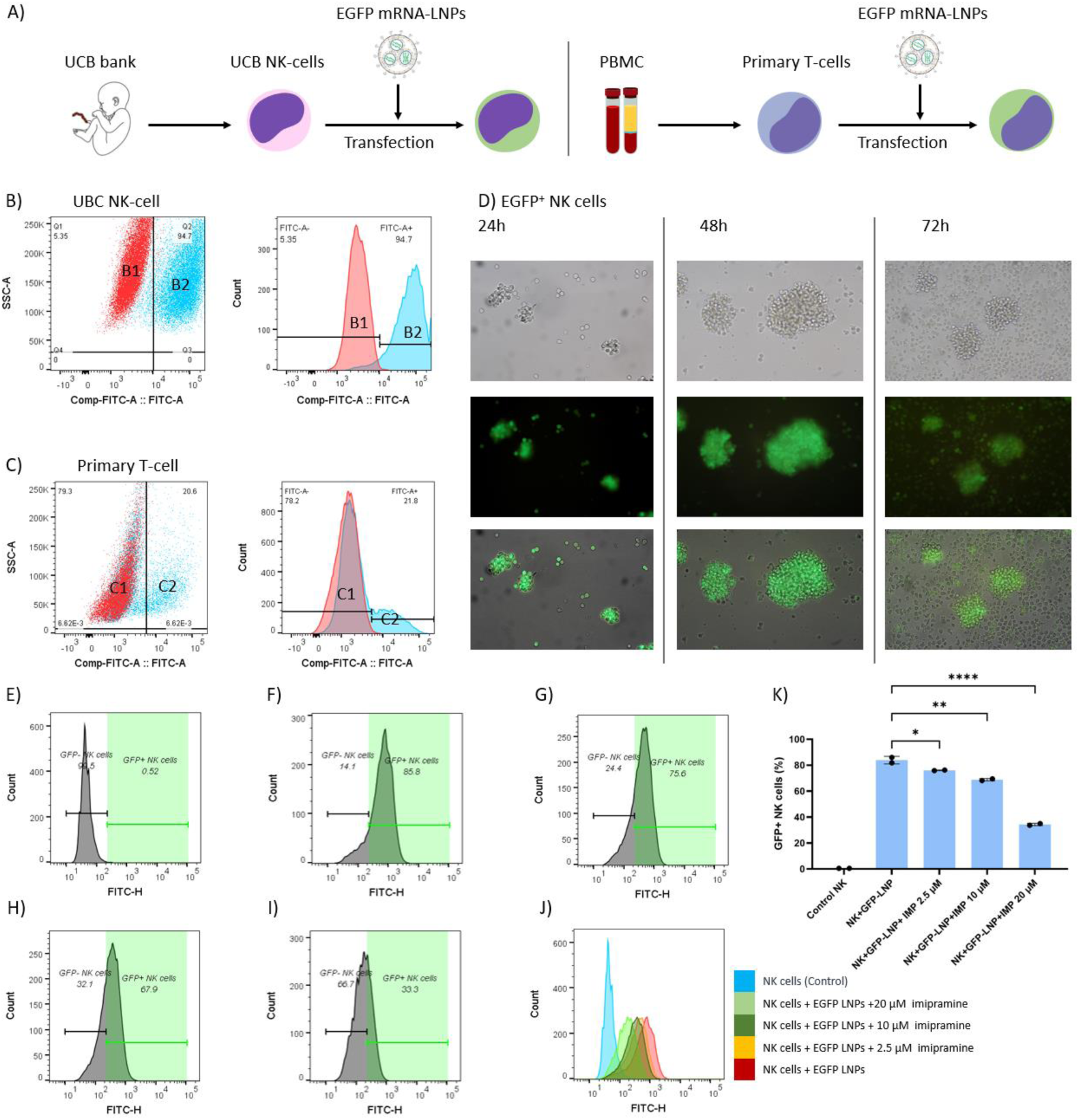
Transfection efficiency of LNPs in UCB-derived NK cells and activated primary T cells. **A**) Schematic representation of EGFP mRNA-LNP transfection in UBC-NK and primary T cells. Transfection efficiency of LNPs encapsulating EGFP mRNA on UCB derived NK cells **(B)** and PBMC derived T cells **(C)** using flow cytometry 24-hours post-transfection. **B1)** Control un-transfected NK cell population in red. **B2)** EGFP^+^ NK cell population and its transfection efficiency in blue. **C1)** un-transfected T cell population in red. **C2)** EGFP^+^ T cell population in blue and its transfection efficiency. **D)** EGFP+ NK cells images at 24-, 48– and 72-hours post-transfection with EGFP-LNPs. Effect of imipramine (IMP) on NK cell mRNA-LNP transfection as a specific macropinocytosis inhibitor. **E**) Control Untransfected NK cells in flow cytometry **F)** NK cells transfected with EGFP mRNA-LNPs. NK cells transfected with EGFP mRNA-LNP in 2.5mM μM (**G),** 10 μM (H), and 20 μM (I) Imipramine. **J)** Overlaid histograms of E-I groups. **K)** Overall Effect of imipramine on NK cell mRNA-LNP transfection.

To evaluate T cell transfection efficiency, we isolated T cells from the PBMCs of healthy donors and activated them using anti-CD2/CD3/CD28 monoclonal antibody beads along with IL-2 in the culture medium. We hypothesized that if LNPs demonstrate efficient transfection in NK cells, they may also exhibit suitable transfection efficiency in T cells. However, our findings indicated that our LNPs were unable to transfect T cells as efficiently as NK cells. The transfection efficiency in activated T cells was observed to be 21.8% at 24-hours post-transfection (Fig.2 C). In conclusion, our LNPs showed superior transfection efficiency in NK cells compared with T cells.

### 3.2. Possible role of macropinocytosis in NK cell mRNA-LNP transfection

To understand the basics of LNP transfection in NK cells, we considered modulating the rate of an endocytic mechanism, macropinocytosis, in NK cells [55]. To do this, we added imipramine, a novel, specific macropinocytosis inhibitor [56], to the NK cell culture medium at varying concentrations 1-hour prior to mRNA-LNP transfection. Our results indicated a significant decrease in transfection efficiency, from 85.8% in the control group to 67.9% and 33.3% when 10 μM and 20 μM of Imipramine were used, respectively (Fig.2 E-K). These results indicate that a reduction in the rate of macropinocytosis is associated with lower transfection efficiencies, suggesting that macropinocytosis may serve as a key endocytic mechanism for mRNA-LNP uptake in NK cells.

### 3.3. Cytotoxicity of anti-CD19 CAR-NK cell against EGFP^+^Raji cell line

Raji cells were used as target cells to evaluate the cytotoxicity of anti-CD19 CAR-NK cells. Raji cells are CD19^+^ and are known to be naturally resistant to NK cells due to the high expression of human leukocyte antigens (HLA) class I molecules on their surface. HLA class I molecules interact with Killer Immunoglobulin-like Receptors (KIR) and natural killer group 2 member A (NKG2A) receptors on NK cells as inhibitory receptors [57, 58]. However, NK cells carrying the anti-CD19 CAR Single-chain variable fragment (scFv) on their surface can effectively recognize Raji cells and mount a strong response against them.

We co-cultured NK and CAR-NK cells with EGFP^+^Raji cells at effector-to-target (E: T) ratios of 5:1 and 1:1. EGFP^+^Raji cells were employed as target cells (Fig.3 A). Propidium iodide (PI) was utilized to label late apoptotic and dead cells within the FITC^+^ population. The results indicated that although nearly all EGFP^+^Raji cells in the control group remained viable after 10–12-hour treatment, both the untransfected NK cell and CAR-NK cell groups demonstrated dose-dependent cytotoxicity in the flow cytometry analysis of EGFP^+^Raji cells. Specifically, we observed 14% and 17% cytotoxicity in the 1:1 and 5:1 untransfected NK cell groups, compared with 35% and 60% cytotoxicity in the 1:1 and 5:1 CAR-NK cell groups, respectively. CAR-NK cell groups demonstrated significantly higher cytotoxicity compared with NK cells, representing the role of CAR in enhanced cytotoxicity of CAR-NK cells (Fig.3 B).

**Fig. 3:**
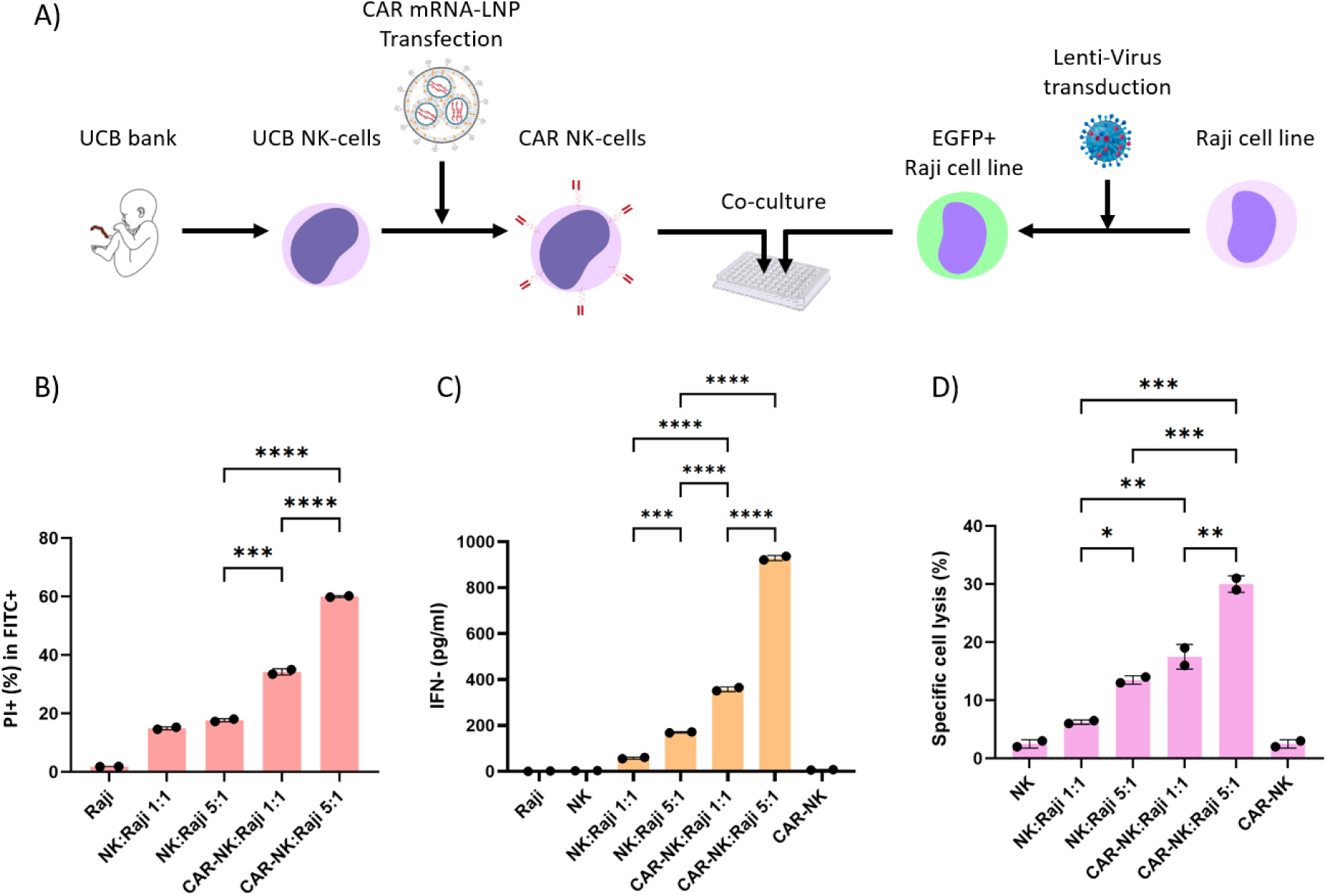
Evaluation of Anti-CD19 CAR-NK Cells cytotoxicity against EGFP^+^Raji cell line. **A)** Schematic representation of CAR-NK and NK cells co-cultured with EGFP^+^Raji cell line. CAR-NK cell production and co-culture with EGFP^+^Raji cell line. Functional assays of CAR-NK and NK cell groups against EGFP^+^Raji cell line in 1:1 and 5:1 E:T ratios. **B)** Comparison of cytotoxicity of anti-CD19 CAR-NK cells and NK cell groups against EGFP^+^Raji cell line using Percentage of PI^+^ cells in FITC^+^ target cell population by flow cytometry in 1:1 and 5:1 E:T ratios. Comparison of IFN-γ Secretion **(C)** and specific cell lysis **(D)** measured by LDH concentrations in NK and CAR-NK groups against EGFP^+^Raji cell line in 1:1 and 5:1 E:T ratios.

To evaluate the cytotoxicity of our CAR-NK cells in more detail, we performed Interferon-gamma (IFN-γ) secretion and Lactate Dehydrogenase (LDH) assays. IFN-γ secretion test is a critical assay for assessing the cytotoxicity of immune cells, especially in T and NK cell therapies. It measures the production of IFN-γ, a cytokine that indicates NK and T cell activation and correlates with enhanced cytolytic function against tumor cells. Thus, elevated IFN-γ levels suggest effective immune responses [59, 60]. On the other hand, LDH assay evaluates cytotoxicity by quantifying the release of lactate dehydrogenase from lysed cells, providing a direct measure of CAR NK and NK cells cytotoxic effect on tumor cells by calculating specific cell lysis. Together, these assays enable us to comprehensively assess the functional capacity of CAR NK and NK cells [61].

We evaluated IFN-γ secretion and LDH concentration using the enzyme-linked immunosorbent assay (ELISA) technique on supernatants of co-culture wells with equal final volumes. The CAR-NK cell group exhibited signifacntly higher IFN-γ secretion and specific cell lysis compared with untransfected NK groups in a dose dependent manner (Fig.3 C, D(confirming the cytolytic potential of CAR-NK cells against Raji target cells.

### 3.4. Cytotoxicity of anti-CD19 CAR-NK cell against primary cells from B-ALL patients

Primary B-ALL cells were isolated from three patients with refractory/relapsed B-ALL during the relapse phase, characterized by a high percentage of blasts in peripheral blood (Fig.4 A). Primary cells were co-cultured with NK cells and CAR-NK cells 24-hours post-transfection in 1:1 and 5:1 E:T ratios. Cytotoxicity of NK cells and CAR-NK cells was measured using LDH for specific cell lysis and IFN-γ secretion test (Fig.4 B, C). LDH and IFN-γ secretion tests show a significant higher cytotoxicity in CAR-NK groups versus untransfected NK cells against Primary B-ALL cells (blast groups) in a dose dependent manner.

**Fig 4.**
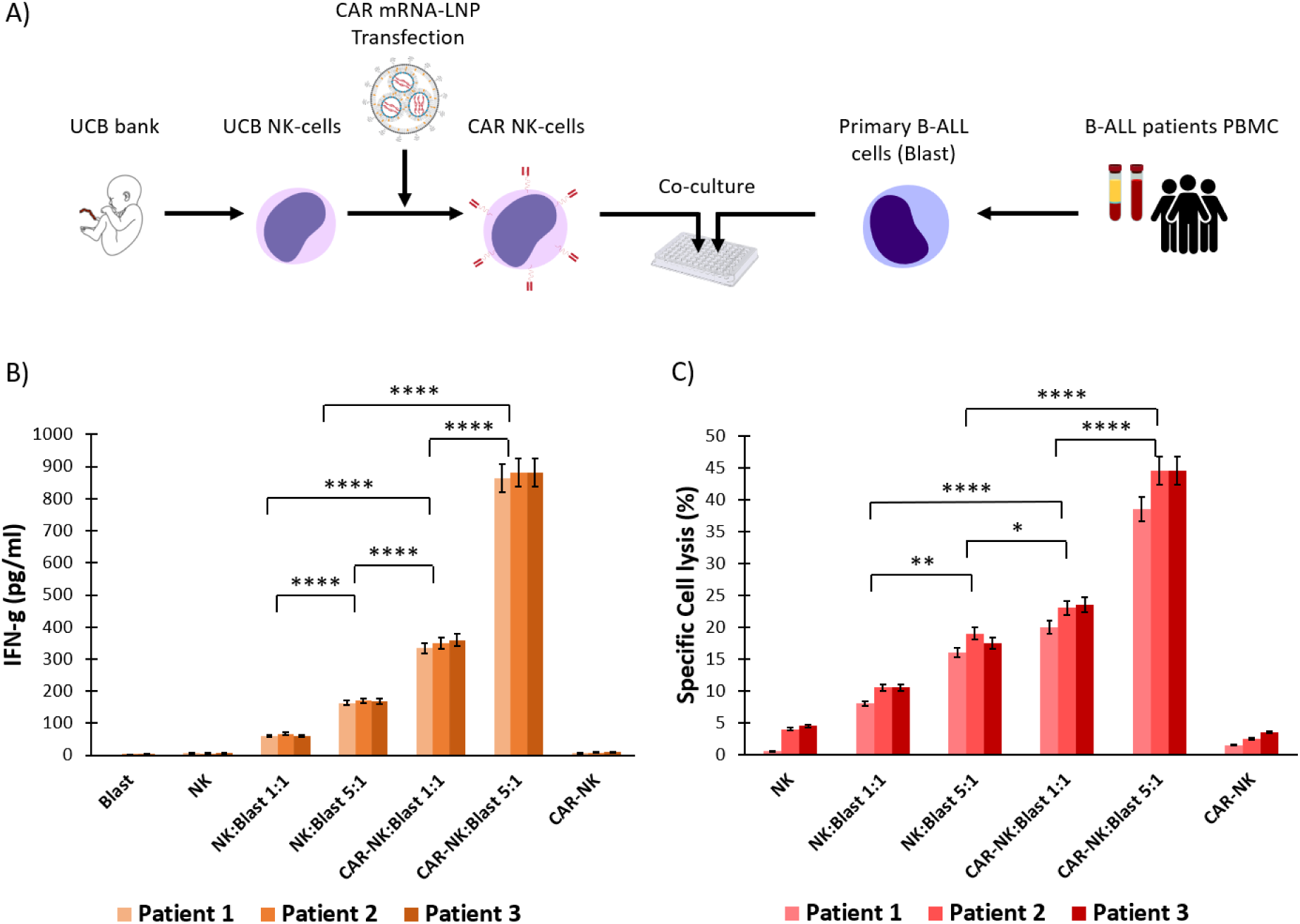
Anti-CD19 CAR-NK cells cytotoxicity against B-ALL patients’ primary cells. **A)** Schematic representation of CAR-NK and NK cells co-cultured with primary B-ALL cells. **B)** Cytotoxicity of NK and CAR-NK cells against primary B-ALL cells by IFN-γ secretion test in 1:1 and 5:1 E:T ratios. **C)** Specific cell lysis measured by LDH concentrations in NK and CAR-NK cell groups against primary B-ALL cells in 1:1 and 5:1 E:T ratios.

## 4. Discussion

NK and T cells play complementary roles in the immune response, exhibiting cytotoxicity against cancer cells [62, 63]. NK cells are key players in the innate immune system, capable of identifying and destroying tumor and virus-infected cells without prior sensitization. In contrast, T cells require activation by antigen-presenting cells (APCs) to recognize specific antigens, playing a vital role in adaptive immunity by generating targeted responses to pathogens and forming immunological memory. [62, 63].

Enhancing the targeting efficiency and functionality of NK and T cells through CAR expression has become a rapidly advancing area in cancer therapy. NK and CAR-NK cells are homogeneous populations that lack T cell receptors (TCRs) and rely on non-MHC-restricted mechanisms for their cytotoxic effects. In contrast, T and CAR-T cells are heterogeneous populations that express TCRs and utilize MHC class I/II-restricted mechanisms for cytotoxicity. This distinction contributes to the “off-the-shelf” potential of CAR-NK cell therapies, while CAR-T cell therapies typically require autologous sources [62, 63], and modifications are required to make “off-the-shelf” CAR-T cell therapies [62, 63]. Furthermore, CAR-NK cell therapy presents reduced risks of graft-versus-host disease (GvHD) and cytokine release syndrome (CRS) in clinical settings, making it a safer alternative to CAR-T cell therapy. Collectively, these characteristics may position CAR-NK cell therapy as a promising and dynamic field in cancer treatment.

To generate CAR-T or CAR-NK cells in an ex vivo setting, both viral vectors for stable genomic CAR integration and mRNA-based approaches for transient CAR expression could be utilized. Transient expression of CAR eliminates the potential risks associated with permanent genetic modifications, such as insertional mutagenesis and oncogenicity in viral vectors. Among mRNA-based approaches, the production of CAR-modified T cells and NK cells using an mRNA-LNP platform is a promising and ongoing field of research [64–66]. This approach has gained significant attention due to its versatility in combination therapy, allowing for the co-encapsulation of a diverse set of siRNAs and therapeutic mRNAs in LNPs. Additionally, delivery efficiency and targeting of LNPs could be enhanced by altering the lipid composition or surface modification, particularly through the decoration of LNPs with diverse types of antibodies, making them applicable for various applications, such as cancer cell therapy, as well as treatment of some autoimmune disorders, and even infectious diseases [67–69]. Furthermore, the mRNA-LNP platform offers a flexible methodology for rapid design and production of CAR constructs, allowing parallelized evaluation and application of various costimulatory domains for efficient targeting of different tumor antigens. This adaptability enables the development of personalized immunotherapies tailored to individual patient needs and tumor characteristics [65, 70, 71].

While T and NK cells were traditionally among hard-to-transfect immune cells, new advancements in LNP formulation have paved the way for efficient mRNA delivery into T, NK and other immune cells [72–75]. For example, in a recent study by Michael J. Mitchell et al., a new ionizable lipid, called C14-4, was developed to serves as a key component in the formulation of LNPs for efficient mRNA delivery in T cells. Using C14-4 LNPs they could effectively deliver CAR mRNA and generate functional CAR T cells which exhibited potent anti-tumor activity comparable to traditional electroporation methods [64, 66, 76]. Regarding NK cells, in a study by Douka et al., novel ionizable lipids were developed for ex vivo NK cell transfection, with “Lipid 5” being optimized to promote transfection efficiency from 36% to 90%. In comparison, the FDA-approved ionizable lipid SM-102 containing LNPs achieved a stable transfection efficiency of 82% in NK cell line (KHYG-1) [48]. Another study also reported that SM-102 containing LNPs could effectively deliver CAR mRNA to PBMC-derived NK cells [49]. Additionally, Nakamura et al., reported nearly 100% transfection efficiency in the NK-92 cell line using their especial ionizable lipid nanoparticle; CL1H6 (CL1H6-LNP) [47].

In this study, we evaluated the transfection efficiency of our proprietary LNPs in UCB-NK and primary T cells. We used EGFP mRNA-LNPs and demonstrated superior mRNA delivery into UCB-NK cells. Notably, we could reach mRNA transfection efficiencies of 94.7% in UCB NK cells, and 21.8% in primary T-cells using our proprietary LNPs (Fig.2 A-D).

To explore mechanisms associated with high transfection efficiency in UCB-NK cells, we focused on the probable role of a particular mechanism of endocytosis, called macropinocytosis. NK cells employ their own balance of endocytic mechanisms that reflect their unique immune functions. Recent studies indicate that NK cells predominantly utilize macropinocytosis, a mechanism that enables them to internalize larger volumes of extracellular fluid while preserving the surface expression of inhibitory receptors such as CD94/NKG2A [55, 77]. Macropinocytosis in NK cells is crucial for regulating the balance between activating and inhibitory signals, enabling NK cells to effectively respond to infected or malignant cells [77]. We hypothesized that modulation of macropinocytosis rate in NK cells may influence the efficiency of LNP transfection by facilitating the uptake of mRNA-LNPs. Imipramine, has been previously shown as a specific macropinocytosis inhibitor [56]. We compared the efficiency of mRNA-LNP transfection in the presence or absence of imipramine (Fig2. E-K), and observed that macropinocytosis inhibition reduced transfection efficiency in NK cells in a dose-dependent manner. Macropinocytosis, thus, may act as a crucial step in mRNA-LNP transfection into NK cells. However, further investigation is required to elucidate its possible implications in other cell types, such as T-cells.

Expanding on these findings, we utilized UCB NK cells for advancing CAR-NK cell production and assessing cytotoxicity against EGFP^+^Raji cell line, representing B-cell malignancies, particularly NHL. Raji cells exhibit a high expression profile of MHC class I molecules, which can bind to inhibitory receptors on NK cells, including KIR and NKG2A, contributing to their resistance against NK cell-mediated cytotoxicity [78–81]. We produced NK cells harboring the second-generation CAR, consisting of anti-CD19 CAR scFv with CD28 and CD3ζ costimulatory domains to target CD19 as a pan B-cell marker. The Anti-CD19 CAR-NK cells generated with our mRNA-LNP platform, showed superior cytotoxicity against EGFP^+^Raji cell line compared with untransfected NK cells (Fig.2). To activate both NK and CAR-NK cells, we supplemented cultures with IL-2 and IL-15 cytokines, which likely contributed to the enhanced cytotoxic responses observed against the target cells. Following these promising results, we extended our investigation to evaluate cytotoxicity of CAR-NK cells against primary B-ALL cells derived from refractory/relapsed adult B-ALL patients. Using the same co-culture protocol as with the EGFP+ Raji cells, we achieved effective cytotoxicity against primary B-ALL cells as well (Fig.4).

At this point we have developed a platform for generating CAR NK cells effective against cancer cells. However, recent advances in the field have the potential to further optimize this platform to ensure the best outcomes in clinical settings. First of all, the stability of mRNA is a critical factor influencing its efficacy in therapeutic applications, particularly in the context of CAR mRNA used for engineering immune cells [82]. The mRNA is inherently prone to degradation due to its linear structure and susceptibility to nucleases, which can lead to diminished CAR expression and reduced therapeutic effectiveness. Using newer generations of therapeutic mRNA with higher stability indices, including circular RNA (circRNA) and self-amplifying mRNA (samRNA) [83–86], may help increasing the time period in which a given mRNA is expressed in the cells. circRNA possesses a covalently closed loop structure that confers increased resistance to degradation by exonucleases, thereby enhancing its stability and translational efficiency. samRNA, on the other hand, relies on its amplification capacity to produce multiple copies from a single mRNA molecule within host cells, thus significantly boosting the expression levels of CAR on cell surface improving the overall therapeutic potential of ex vivo engineered immune cells [85, 86].

Additionally, further optimizations of the CAR construct itself could enhance the efficacy and functionality of CAR-NK cells. For example, incorporating an IL-15 domain into the CAR construct can further enhance the therapeutic effectiveness. IL-15 enhances NK cell activation and survival while uniquely avoiding the expansion of regulatory T cells, making it particularly advantageous for therapeutic applications. The inclusion of IL-15 in culture systems notably improves the expansion and persistence of CAR-NK cells, thereby enhancing their efficacy against tumors [87, 88].

In this study, we harnessed the potential of the mRNA-LNP platform for CAR armored cell therapy applications. We could achieve efficient CAR mRNA transfection efficiencies in UCB NK cells and, to a lesser extent, in primary activated T cells. Our investigation into the impact of macropinocytosis on NK cell transfection efficiency highlights the probable role of macropinocytosis, and potentially other endocytic mechanisms, in the transfection of NK and T cells by LNPs. The anti-CD19 CAR NK cells developed using our mRNA-based approach exhibited effective cytotoxicity against both the EGFP^+^Raji cell line and primary B-ALL cells derived from three B-ALL patients. These promising results lay a solid groundwork for future preclinical and clinical studies.

## 5. Materials and Methods

### 5.1. Preparation and culture of required cells

To generate a stable Raji cell line expressing EGFP, HEK-293 cells were co-transfected with psPAX2 (Addgene plasmid #12260), PLX_307 GFP (Addgene plasmid #184492) and pMD2.G (Addgene plasmid #12259) using Lipofectamine (Thermo). Virus-containing supernatants were collected at 48– and 72-hours post-transfection and filtered through 0.45 μm filters to remove cellular debris. Raji cells were then transduced with the harvested lentivirus in the presence of Polybrene (10 μg/mL) to enhance transduction efficiency. EGFP-expressing cells were selected using puromycin (10 μg/mL).

Primary B-ALL cells from 3 refractory/relapsed adult B-ALL patients were isolated in relapse phase with high %blast in peripheral blood and were used for in vitro studies of CAR-NK cytotoxicity. The blood samples were collected following informed consent (IR.TUMS.HORCSCT.REC.1403.014) and PBMCs were collected by a density-gradient technique and harvested for following cytotoxicity tests.

PBMCs were collected by a density-gradient technique (Ficoll-Histopaque; Sigma) from healthy donors following informed consent and harvested for T cell isolation. CD3^+^T cells, were isolated and selected from the PBMC with T cell isolation kit, (Supp Fig.1) (Miltenyi Biotec, Inc., San Diego, CA) and were stimulated with anti-CD2/CD3/CD28 monoclonal Ab beads (Dynabeads, Milteny Biotec, Inc., San Diego, CA) at 1:1 ratio (bead: cell). Recombinant IL-2 was added at a final concentration of 100 IU/ml and incubated for 72-hours. Then Activated T cells were used for transfection efficiency of LNPs.

CD3-CD56^+^ NK cells were isolated from UCB units provided by the Shariati Hospital UCB Bank and were isolated by a density-gradient technique (Ficoll-Histopaque; Sigma). CD3-CD56+ NK cells, purified with an NK isolation kit (Miltenyi Biotec, Inc., San Diego, CA) (Supp Fig.1), were stimulated with irradiated (100 Gy) EBV-LCL (20:1 feeder cell:NK ratio) and recombinant human IL-2 (500 U/mL; Miltenyi Biotec, Inc., San Diego, CA) and IL-15(10ng/ml; Miltenyi Biotec, Inc., San Diego, CA) and 10% FBS in RPMI 1640 (Biosera, France) on day 0. NK cells were transfected with mRNA-LNP on day +7 in upright culture flask for 24h and then CAR-transfected NK cells were used for further functional assays.

EBV-LLCL feeder cells were cultured in an RPMI-1640 medium supplemented with penicillin/ streptomycin and 10% FBS.

### 5.2. EGFP and anti-CD19 CAR mRNA Production

To prepare our in vitro transcription (IVT) plasmids, EGFP and anti-CD19 CAR sequences were amplified from EGFP (Addgene plasmid # 176015) and pSLCAR-CD19-28z (Addgene plasmid # 135991) plasmids and cloned into proprietary IVT plasmid backbone by Gibson cloning. The IVT plasmids contained EGFP and a second-generation anti-CD19 CAR with CD28 and CD3ζ costimulatory domains, respectively. These plasmids were then digested and linearized using restriction enzymes, which was confirmed by gel electrophoresis. IVT reactions were performed on the linearized plasmids containing the T7 promoter, generating EGFP and anti-CD19 CAR mRNAs. The mRNA was co-transcriptionally capped with CleanCap reagent (TriLink #N-7413) to enhance its stability and translation efficiency. Finally, the mRNA was purified using the Monarch RNA Cleanup Kit (NEB #T2030) following the manufacturer’s protocol, and its concentration was measured using a NanoDrop spectrophotometer.

### 5.3. Formulation of EGFP and anti-CD19 CAR mRNA

Two solutions were prepared with lipid mix (DSPC, DMG-PEG2000, Cholesterol, Ionizable lipid with molar ratio of 10:1.5:38.5:50) in extra pure ethanol and Anti-CD19 CAR and EGFP mRNA in citrate buffer (50 mM). mRNA and lipid solutions were mixed at the ratio of 3:1 (aqueous: organic phase) within a dedicated microfluidic device to handle encapsulation. LNPs containing mRNA were then purified and buffer exchanged using Amicon® Ultra-15 centrifugal filter units (30–100 kDa). The size and PDI of the nanoparticles were measured by dynamic light scattering (DLS) (ZetaSizer Ultra, Malvern USA), and the encapsulation of mRNA within the LNPs was evaluated using a gel retardation assay.

### 5.4. Transfection of mRNA-LNP to NK and T Cells

To assess the transfection efficiency of LNPs, the EGFP mRNA-LNP was used to transfect NK cells and T cells. NK cells were transfected with 1-2μg EGFP mRNA-LNP per 10^6^ cells on day +7 in upright culture flask at 37°C in a humidified incubator with 5% of CO2 and then harvested for flowcytometry analysis with BD FACSLyric system (BD Biosciences). Optimal transfection conditions were determined using EGFP mRNA-LNP expression results, and then applied to NK cells transfection with the CAR mRNA-LNP. The resulting CAR-NK cells were harvested for further functional assays. To evaluate the transfection efficiency for T cells, CD3^+^ activated T cells with anti-CD2/CD3/CD28 mAb beads and IL-2 were used with the same protocol as NK cells and a dosage of 1-2μg EGFP mRNA-LNP per 10^6^ T cells.

### 5.5. Evaluation of cytotoxicity of CAR-NK cells against EGFP^+^Raji Cell Line and Primary cells

NK cells and CAR-NK cells (effector cells) were co-cultured with EGFP^+^Raji and Primary cells at the appropriate E:T ratios (1:1 and 5:1) in round-bottom 96-well plates at 37 °C in a humidified incubator with 5% of CO2 for 6 h. Propidium iodide (PI) (Sigma-Aldrich, St. Louis, MO) was utilized at a concentration of 50 μg/ml for the labeling of DNA in dead cells, which were subsequently analyzed by flow cytometry. Primary B-ALL cells were labeled using CSFE dye prior to Co-culture with NK and CAR-NK cells.

Co-cultured cells were harvested, washed twice with phosphate-buffered saline (PBS), and resuspended in flow cytometry staining buffer before adding PI staining solution, followed by a 1-minute incubation in the dark prior to flow cytometric analysis to distinguish live from dead cells. EGFP^+^Raji and CSFE-labeled primary B-ALL cells were gated as target cells in FITC channel and then measured %PI^+^ cells in each group.

IFN-γ secretion was quantified using the Human IFN-γ Quantikine ELISA Kit (Catalog # DIF50C) from R&D Systems, designed to measure human IFN-γ levels in cell culture supernatants. The assay was performed according to the manufacturer’s instructions, ensuring accurate and reliable results. After 10-12 hours of incubation, the LDH assay was done according to manufacturer’s guidelines (Roche, Switzerland). NK/CAR-NK cytotoxicity was analyzed as Target lysis (%) = [(effector and target cell mix – effector cell control) – low control] / (high control – low control) * 100.

### 5.6. Statistical Analysis

Statistical analysis was performed using a t-test for comparisons between two groups. For comparisons involving three or more groups, one-way ANOVA was applied. Results were deemed statistically significant at p < 0.05, with significance levels denoted as follows: *P < 0.05, **P < 0.01, ***P < 0.001, and ****P < 0.0001. To assess overall differences among patient population groups, two-way ANOVA was used followed by Tukey’s multiple comparisons test. Graphs were generated using GraphPad Prism software, version 9.0.0, and flow cytometry analysis was performed using FlowJo software, version 10.

## 6. Data availability

All data generated or analyzed during this study are included in this published article.

## 7. Ethics statement

All volunteers were provided with written informed consent and the protocol was reviewed and approved by the institutional review board of Tehran University of Medical Sciences.

## 8. Authors’ contributions

The authors V.K., M.V., S.D., and Mo.A. designed the study. R.A. and V.K. designed the genetic constructs. M.S.M. with the help of Ma.A. and H.S.S. prepared mRNA products. M.S., S.D., and V.K. designed the LNP formulation. M.S. with the help of H.S.S. conducted the mRNA-LNP formulation. Mo.A. and M.V. provided NK and T cells. Mo.A., S.T. and H.S.S. conducted transfection efficiency tests. Mo.A., S.T. and H.S.S. conducted functional assays. M.B., M.O.A helped in patient recruitment. H.S.S. wrote the manuscript draft under the supervision of V.K. and Mo.A.. H.S.S. produced the figures. A.M., Mo.A., M.S., S.D, and V.K. edited the manuscript.

## 9. Acknowledgments

We would like to express our sincere gratitude to Cell Therapy and Hematopoietic Stem Cell Transplantation Research Center in Shariati Hospital and Tehran University of Medical Sciences for their unwavering support throughout this project, which was instrumental in facilitating its successful completion. We would also like to thank Seyed Hossein Kiaie, Ehsan Ansari and Seyed Milad Safar Sajadi as former ReNAP members, which greatly contributed to the success of this study.

## 10. Competing interests

The authors V.K. and S.D. are management board member and employees at ReNAP. M.S., Ma.A., A.M., and M.S.M. are current employees at ReNAP. R.A. is former employee at ReNAP. Mo.A. is Associate Professor of Laboratory Hematology and Blood Banking; M.V. is head of research institute and Associate Professor of hematology and oncology; M.B. is assistant professor of hematology and oncology; and S.T. is PHD student at Hematopoietic Stem Cell Transplantation Research Center in Shariati Hospital, Tehran university of medical sciences. H.S.S. is medical student at medical science department of Tehran university of medical sciences. The authors declare no other relationships or activities that could appear to have influenced the submitted work.

