## Supplemental data for "Effective targeting of CD19 positive primary B-ALL cells using CAR-NK cells generated with mRNA-LNPs"

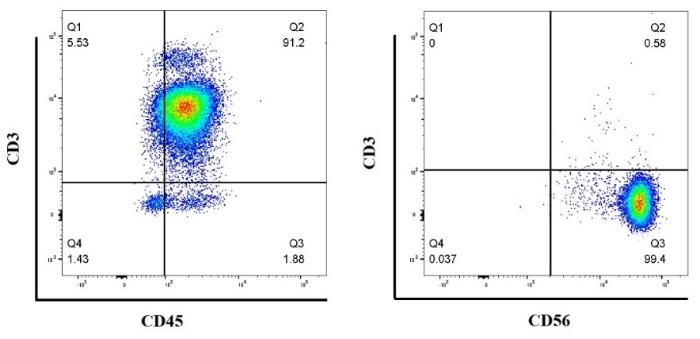

**Supp Fig. 1:** Flow cytometry of A) CD3^+^CD45^+^ T-cells isolated from PBMC of healthy donors, B) CD3^-^CD56^+^ NK cells derived from UCB.

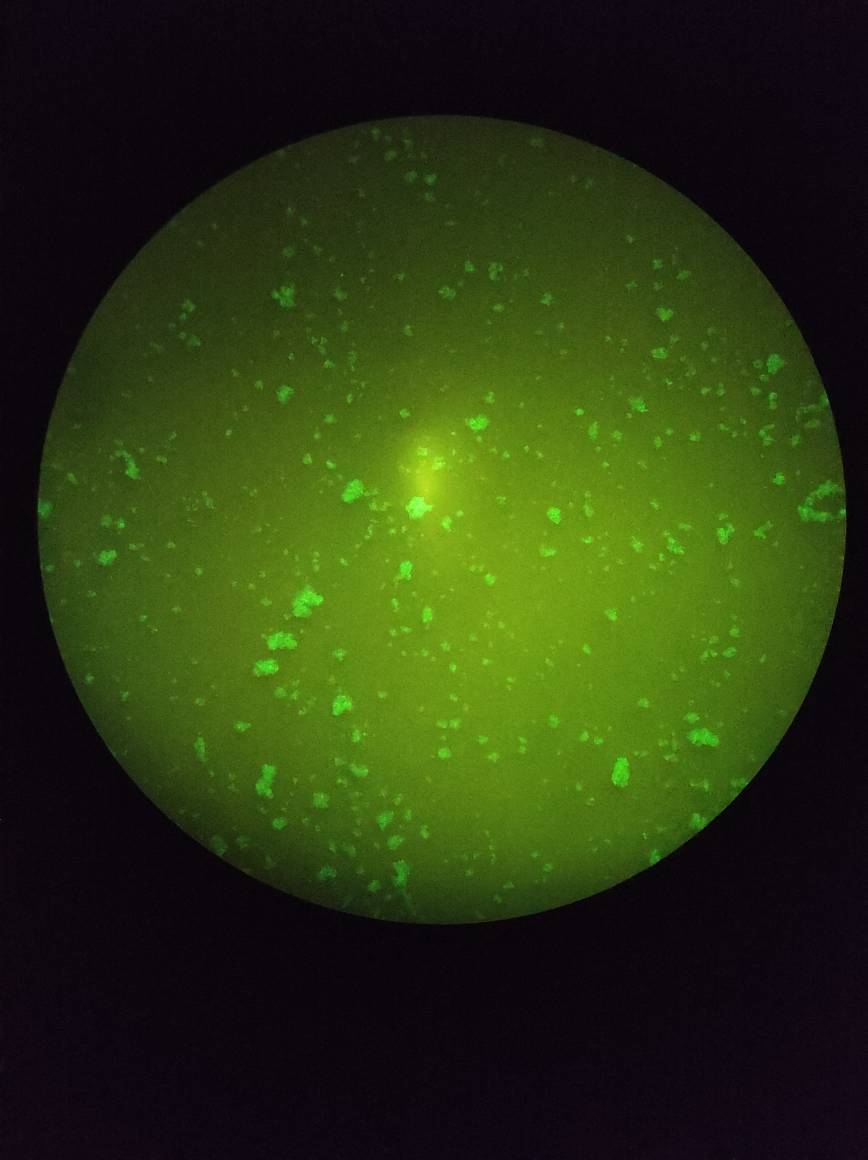

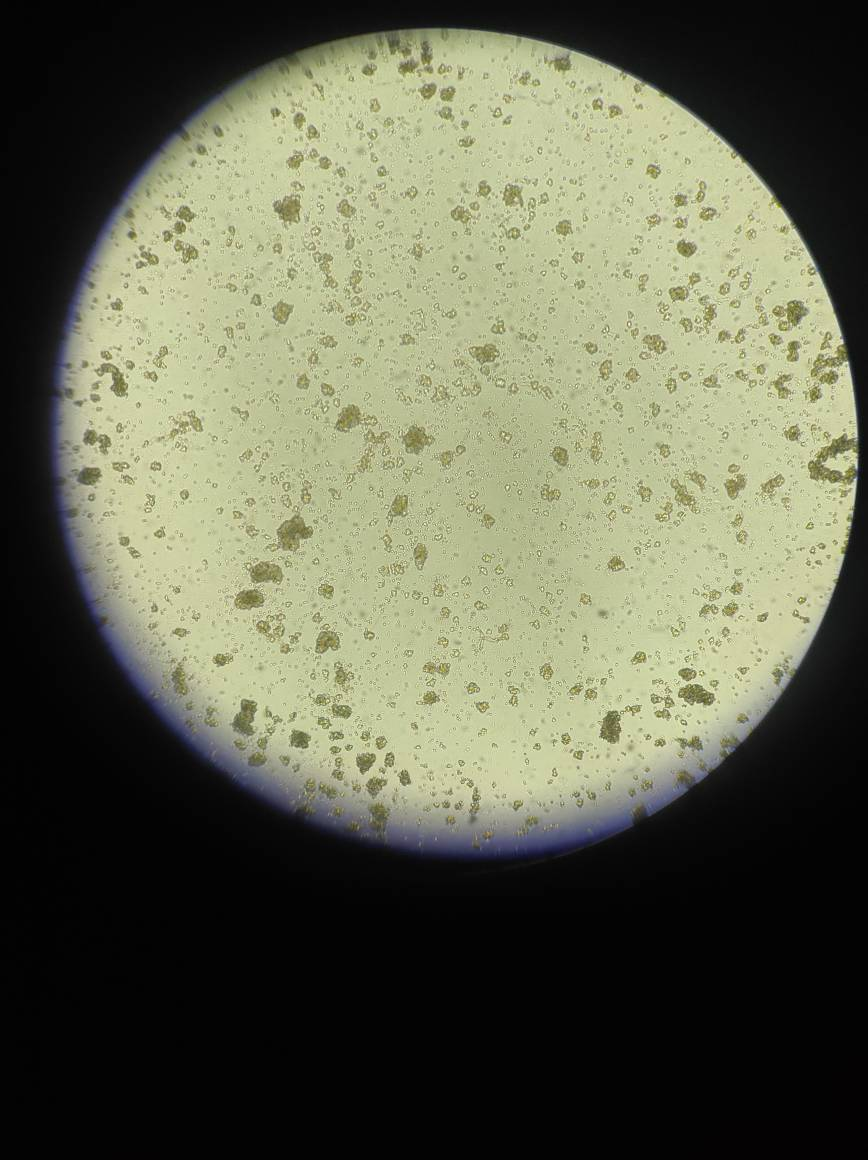

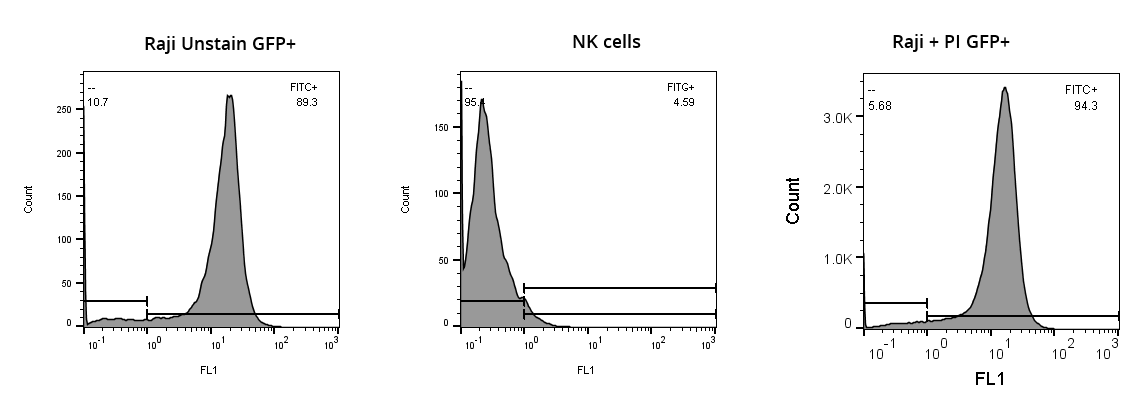

**Supp Fig.2:** Flow cytometry and microscopy of stable EGFP^+^Raji cell line.
